# Investigating the dynamics of the aquatic community in Oslofjord through time series analysis of eDNA

**DOI:** 10.1101/2025.02.05.636659

**Authors:** Sibusiso Mahlangu, Isabelle Ewers, Cintia Oliveira Carvalho, Audun Schrøder-Nielsen, Micah Dunthorn, Hans Erik Karlsen, Grete Sørnes, Louise Chavarie, Dag Endresen, Jonathan Stuart Ready, Hugo J de Boer, Quentin Mauvisseau

## Abstract

Understanding temporal dynamics of marine communities is critical for assessing ecosystem health and guiding conservation efforts. Here, we conducted a survey using environmental DNA (eDNA) metabarcoding with two primer sets, MiFish and Elas02, to investigate seasonal and interannual changes in the fish community of Oslofjord (Norway) over two consecutive years. Using the MitoFish reference database through the GBIF querying tool, we identified 61 fish species and found significant changes in the dominant taxa between seasons and years. *Clupea harengus* (Atlantic herring) consistently peaked in early spring, while *Scomber scombrus* (Atlantic mackerel) dominated winter months in the second year. The MiFish primer set showed increased species richness in the second year, whereas the Elas02 primer set showed stable richness despite compositional turnover. Non-metric multidimensional scaling (NMDS) revealed distinct community separation between years in presence/absence data, driven by species turnover rather than abundance read changes. Our findings support the use of eDNA metabarcoding to capture fine-scale temporal dynamics and emphasise the importance of multi-year datasets for distinguishing ecological trends from stochastic changes. This strategy improves monitoring practices for marine ecosystems under anthropogenic stressors.

## Introduction

Anthropogenic activities, including pollution (Jaureguiberry et al., 2022; Pawar, 2016), introduction of alien species (Bax et al., 2003; Katsanevakis et al., 2013), habitat degradation(Prakash & Verma, 2022; Stuart-Smith et al., 2021), and over-exploitation of resources (Nikolaou & Katsanevakis, 2023; Worm et al., 2021) are leading to the loss of essential marine species, which causes disturbances in food web dynamics and has cascading effects (Fung et al., 2015), including poor nutrient cycling (B-Béres et al., 2023; Sanz-Lázaro et al., 2021), proliferation of invasive species (De-la-Torre et al., 2023; Galià-Camps et al., 2024), and biogeographical shifts (Frigstad & Ramon, 2024; Porter et al., 2013). Our understanding on how human-induced activities impact marine ecology and biodiversity has been significantly improved by monitoring methods including the capture or observation of specimens and their identification using morpho-taxonomic classification methods to estimate species presence, abundance, and community structure (Keck et al., 2022). However, these approaches have shown limitations such as high costs and logistical difficulties when accessing remote areas (Sevellec et al., 2021), the time and labour required (Gibson et al., 2015), the lack of expertise to accurately identify small or cryptic species (Keck et al., 2022), and the risk of missing elusive species that are difficult to observe (Mathieu et al., 2022; Stoeckle et al., 2020). To mitigate these limitations, molecular-based techniques relying on environmental DNA (eDNA) detection offer an alternative approach for biodiversity assessments and monitoring across large spatio-temporal scales (Courtaillac et al., 2024; Perry et al., 2024; Sales et al., 2021). Effective implementation of management strategies requires reliable approaches for environmental data collection combined with early detection of changes in community composition across various spatial and temporal scales—eDNA can provide the latter (Keck et al., 2022; Wang et al., 2024).

Environmental DNA refers to the intra- and extracellular DNA released by organisms into their surrounding environments and is used as a proxy for detecting species presence (Cindy et al., 2021; Taberlet et al., 2012; Wang et al., 2024). In recent years, eDNA metabarcoding-based biodiversity assessments have gained recognition in marine research and monitoring as a complementary method to morphology-based approaches (Blattner et al., 2021; Djurhuus et al., 2020; Keck et al., 2022; Mathieu et al., 2022; Sevellec et al., 2021; Wang et al., 2024). Coupling eDNA-based monitoring with time-series analysis allows the investigation of fluctuations and changes in community composition (Djurhuus et al., 2020; Sevellec et al., 2021). Time-series analysis is crucial when exploring seasonal cycles of biodiversity and for detecting phenological shifts (Collins et al., 2022; Sander et al., 2024; Sigsgaard et al., 2017). Through the detection of shifts in the community or early invasive species, time-series analysis can function as an early-warning system (Djurhuus et al., 2020). This is therefore essential when interpreting long-term biodiversity patterns to obtain insights into ecosystem stability and resilience or when assessing the long-term impact of stressors such as climate change (Stoeckle et al., 2021) or sporadic events such as algal blooms (Jacobs-Palmer et al., 2021).

To be able to detect and monitor seasonal variations in marine fish communities, it is essential to have a reference database of sequences available that provides species occurrence records (Berry et al., 2021; Lim & Thompson, 2021). Good taxonomic coverage of reference sequences of the targeted organism group and marker gene region for taxonomic assignment are also required (Blackman et al., 2023; Ruppert et al., 2019). The Global Biodiversity Information Facility (GBIF) integrates species occurrence records from scientific publications, museum collections, species monitoring, citizen science efforts, and eDNA studies (Garcia-Rosello et al., 2023; GBIF, 2024). The deposited data is used to analyse species distributions and biodiversity trends for research, conservation, and sustainable management purposes. GBIF provides tools allowing querying taxonomic reference databases specialized for specific markers and organisms (Abarenkov et al., 2024; Frøslev et al., 2024; GBIF, 2024), such as the MitoFish database (Iwasaki et al., 2013; Sato et al., 2018). MitoFish is a comprehensive and standardized fish mitochondrial genome database enabling the taxonomic identification of fish environmental DNA (Iwasaki et al., 2013; Sato et al., 2018). MitoFish provides a curated reference library containing mitochondrial sequence data of more than 45,000 species to enable species-level identification of sequences generated by eDNA metabarcoding, making it possible to track changes in fish community composition over time (Zhu et al., 2023).

The Oslofjord is under various pressures associated with long term human activities, and while ecological conditions have improved in some parts of the fjord, new invasive species are spreading and an increasing proportion of species, such as cod (*Gadus morhua*) are disappearing or currently endangered (Frigstad & Ramon, 2024). To mitigate this, the Norwegian Directorate of Fisheries (Fiskeridirektoratet) released several approaches to promote sustainable management of the marine resources in the fjord, including the introduction of three major non-fishing areas (Fiskeridirektoratet, 2024). However, as climate change is expected to worsen the current situation, a recent report has highlighted the necessity of time series to better understand natural fluctuation and possible population changes in the fjord (Frigstad & Ramon, 2024). Here, we combined eDNA-based detection and time-series analysis to investigate seasonal changes in the marine fish community in the Oslofjord. Following proof of concept studies investigating the benefit and limitation of eDNA-based monitoring in this area (Carvalho et al., 2024; Kvalheim et al., 2024), we now aim to assess if eDNA-based monitoring allows the study of community structure changes over time to better guide conservation and management decisions, and eventually assess the effects of potential actions supporting recovery processes.

## Methods

### Study site

eDNA samples were collected in the Oslofjord near Drøbak (Norway), over two years, from March 2022 to January 2024, across a sampling transect as described in Carvalho et al. (2024). Samples collected from March 2022 to January 2023 are already described in Carvalho et al. (2024), and samples collected from March 2023 to January 2024 are described in (Table S1).

### eDNA sampling

eDNA samples were collected as described in Carvalho et al. (2024). In brief, surface water was collected at each sampling site and three natural replicates were filtered using sterile 0.8 μm Whatman (Cytiva, USA) mixed cellulose ester filters (22 mm diameter) and sterile 22 mm Swinnex (Merck Millipore, Germany) filter holders, using a Vampire Sampler (Bürkle GmbH, Germany) pump system. We collected surface water to allow a reliable comparison with the eDNA surveys performed in (Carvalho et al., 2024, Kvalheim et al., 2024), and while we acknowledge that this could have led to a biased detection of pelagic species compared to benthic ones, the relatively low depth in the fjord by Drøbak (20 m) should mitigate this potential issue. Volume filtered for each natural replicate at each site and sampling event ranged from 400 to 1000 mL depending on turbidity (Carvalho et al., 2024, Table S1). One negative control consisting of 1 L ddH2O was filtered alongside eDNA samples at each sampling event. Following filtration, filters were stored at -20 °C until DNA extraction. Environmental conditions recorded at each site during each sampling event included temperature (°C), conductivity (mS), salinity, total dissolved solids (TDS, g/L), and pH (Carvalho et al., 2024, Table S1). To minimize potential cross-contamination, sterile equipment and disposable gloves were used throughout the sample processing and sampling events. A total of 192 eDNA samples were collected from March 2022 to January 2024 (i.e. 180 eDNA samples and 12 field negative controls).

### eDNA analysis

eDNA samples were extracted using the QIAGEN Blood and Tissue Kit (Qiagen, Netherlands) in a dedicated PCR-free room, following the manufacturer’s instructions with slight modifications as in Carvalho et al. (2024). PCR amplifications were conducted using two primer sets targeting a short fragment of the mitochondrial 12S rRNA region, the MiFish-U-F 5’-GCCGGTAAAACTCGTGCCAGC-3′ and MiFish-U-R 5′-CATAGTGGGGTATCTAATCCCAGTTTG-3′ primer set (Miya et al. 2015) and the Elas02F/R 5′-GTTGGTHAATCTCGTGCCAGC-3′ and 5′-CATAGTAGGGTATCTAATCCTAGTTTG-3′ primer set (Miya et al., 2015; Taberlet et al., 2018). Only eDNA samples collected from March 2023 to January 2024 were processed and amplified in this study, and raw sequencing data using the same primer sets were retrieved for the period spanning from March 2022 to January 2023 from (Carvhalo et al., 2024). All samples were amplified in triplicate, with the inclusion of 5 extraction blanks and 2 NTCs (i.e. Non-Template Control, where eDNA was replaced with ddH2O). All PCR amplifications were conducted using indexed primers as in Fadrosh et al. (2014). PCR were conducted in a final volume of 20 μL using 10 μL of 2X Accustart Toughmix II (QuantaBio, USA), 0,5 μL of each indexed primer (10 μM each), 7 μL of nuclease free water and 2 μL of extracted eDNA. The PCR amplification parameters were as follows: initial denaturation at 94 °C for 3 min, followed by 35 cycles of 94 °C for 20 s, 52 °C for 20 s and 72 °C for 30 s and then a final extension of 72 °C for 3 min. PCR products were then visualised on 1% agarose gels and quantified using the ImageLab Software v6.0 (Bio-Rad Laboratories, USA). To ensure an equimolar representation of the amplified samples, a Biomek 4000 liquid handling robot (Beckman Coulter, USA) was used to merge amplicons into a library. The library was then cleaned using 10% of Illustra ExoProStar (Cytiva, USA) and 1.0X AMpure XP (Beckman Coulter, USA) bead clean. Size selection of the targeted fragments for each primer set was done using a BluePippin (Sage Science, USA). Quality control was performed by visualising the library on a Fragment Analyzer (Agilent, USA) using the High Sensitivity Genomic DNA kit, and both MiFish and Elas02 libraries were sequenced on an Illumina Miseq platform using v2 2*150 bp chemistry (Illumina, USA).

### Bioinformatics

Bioinformatics was conducted together on the raw sequencing files retrieved from (Carvalho et al. (2024) and from the samples sequenced in this study. The Raw Fastq files were demultiplexed and processed for primer removal using cutadapt version 4.4 (Martin, 2011), and reads not matching primers were discarded. DADA2 v1.28.0 (Callahan et al., 2016) was used for denoising and sequence pair construction. Quality score plots were carefully examined, and criteria for Phred score quality, forward and reverse lengths were trimmed when the Phred score dropped below 30. Reads were de-duplicated and forward and reverse error models were generated for nucleotide error estimation. Reads were merged into amplicon sequence variants (ASVs), with chimaeras excluded. All DADA2 arguments were set at default except for the truncating of sequences using the ‘truncLen’ argument of the filterAndTrim function, which depended on the dataset’s quality plots. A custom R function was used to create a FASTA file with all output sequences and corresponding sequence table (ASV table) with the read counts for the corresponding ASV’s in each sample (Salib, 2022). A template code is found alongside the raw data in Zenodo at https://doi.org/10.5281/zenodo.14809757. Each dataset was processed independently (i.e., four datasets, one for each sampling year and primer set). The generated FASTA was blasted using the 12S Sequence ID tool on the Global Biodiversity Information Facility (GBIF) site (gbif.org 2024) to taxonomically classify the resultant ASV reads. This tool queries sequences against the MitoFish v3.97 database, which only contain 12S Fish reference sequences. The resultant taxonomy table was cleaned to retain only fully classified sequences associated with fish, with a 97% similarity score to the reference database, and used together with the ASV table for downstream analysis.

### Data treatment

To remove potential contaminants, false positives, and sequencing errors, we filtered the ASV count table. Firstly, we removed the maximum number of reads detected in the negative or extraction blank control samples (whichever had the maximum read) for each associated ASV across all samples (Carvalho et al., 2024). Secondly, we removed ASV’s that had a low frequency (0.01% relative read abundance) per taxon per library (Milhau et al., 2021). Thirdly, we pooled the PCR replicates and site replicates to reflect the overall eDNA signal rather than technical variations. Finally, reads below 0.01% of the entire dataset read abundance were removed in order to retain species that contribute meaningfully to the overall dataset and have biological relevance (Karstens et al., 2019). This was done for each dataset independently (each year with each primer set).

### Statistical analysis

All statistical analysis and graphics were done using the R software v4.3.2 (RStudio v2024.04.2 +764). To visualise species composition across the months and years, a relative abundance barplot was constructed using ggplot 2 v3.5.1 (Wickham, 2016). Species richness computation between sampling events (months) and years (Year 1 vs Year 2) was estimated using the estimate_richness function from the phyloseq v1.44.0 (McMurdie & Holmes, 2013). We visualised the differences in read abundance across sampling months by producing a bubble plot using the ggplot function (Wickam, 2016). To assess the differences in species composition and abundance by year, we performed non-metric multidimensional scaling (NMDS) analyses using the vegan v2.6.6.1 (Dixon, 2003) using two datasets, one with only presence/absence data (1 = presence, 0 = absence) vegdist R function with Bray-Curtis distance calculation, and the other dataset with rarefied reads using the avgdist R function using the lowest read depth across all samples using Bray-curtis distance (Dixon, 2003). The PERMANOVA analysis from the vegan package adonis2 function (Oksanen & Simpson, 2022) was used to test for differences between the community composition between the two years for each primer.

## Results

A total of 14 314 724 and 13 457 523 raw reads were obtained following metabarcoding amplification and NGS sequencing for Year 1 and Year 2. Following bioinformatic processing, 3 066 584 and 3 335 299 reads were assigned to MiFish and Elas02 respectively for the Year 1, and 1 813 903 and 2 337 568 reads were assigned to MiFish and Elas02 for the Year 2. After data cleaning and retaining reads assigned only to fish, we ended up with 48 345 (mean of 8 057.5 per sampling event, i.e. all replicates and sites combined at a given month) and 435 055 (mean of 72 509.2 per sample event) reads assigned to MiFish and Elas02 respectively for Year 1, and 292 541 (mean of 4875 per sample event) and 223 079 (mean of 37179.83 per sample) reads assigned to MiFish and Elas02 respectively for Year 2. Following taxonomic assignment, we identified a total of 61 fish species found across both sampled years, and primers combined are provided in supplementary file Table S1 and Table S2.

Species composition varied throughout the sampling period for both years using both the MiFish and Elas02 primer sets (Figure 1), with a change in species relative abundance among months and between years using both primer sets (Figure 1). For the MiFish primer set, reads for *Clupea harengus* were dominant in March of both Year 1 and Year 2 (Figure 1), while also being present throughout most months in both years (Figure 2). May of the Year 1 was dominated by C*. harengus* and *Pollachius virens* reads, whereas May of Year 2 was dominated by reads of *Sprattus sprattus*. *Scomber scombrus* dominated July and September in Year 1, while those months were dominated by *Gobiusculus flavescens* reads in Year 2, with November of both years dominated by *G. flavescens* reads (Figure 1). January of Year 1 shows similar read abundance for several species, without any clearly dominant species, while January in Year 2 was dominated by *S. scombrus*.

**Figure 1.**
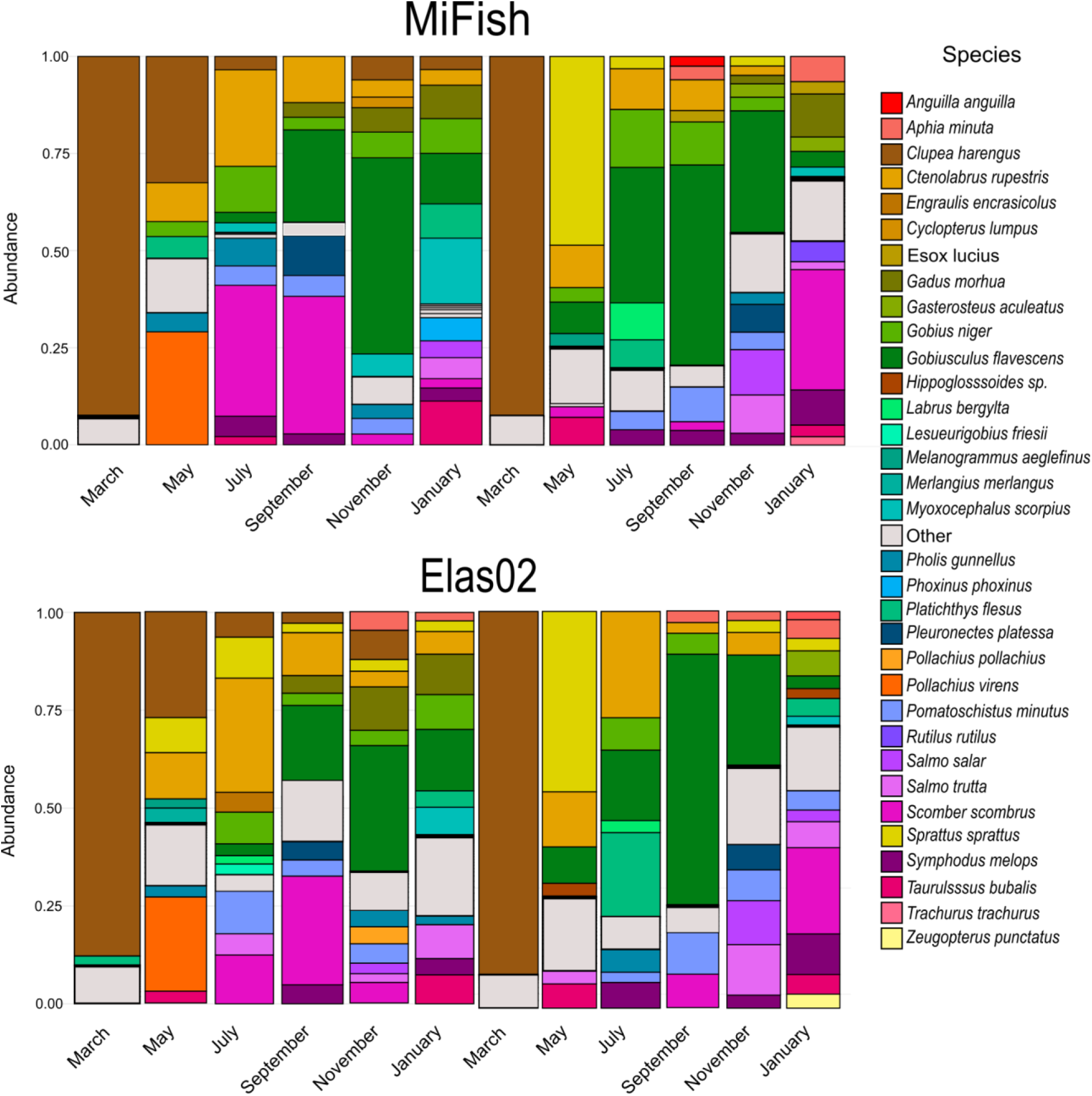
Relative read abundance of the 33 most abundant fish species detected through eDNA metabarcoding using both primer sets every two months during a two-year time period from March 2022 to January 2024.

**Figure 2.**
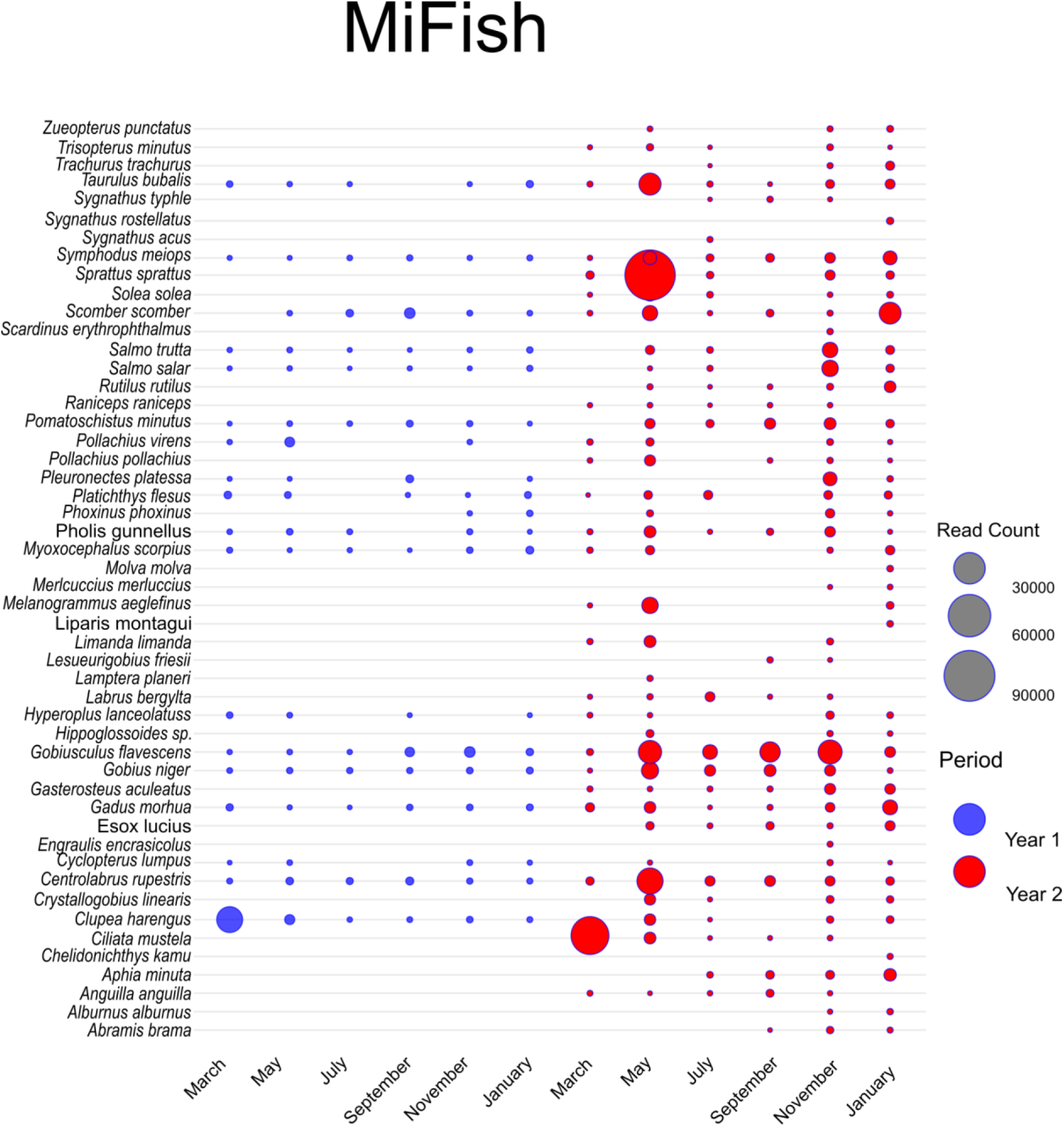
Bubble plot showing the relative read abundance of each detected fish species using the MiFish primer set across all sampling events

For the Elas02 primer set, reads of *C. harengus* were dominant in March of both Year 1 and Year 2 (Figure 1), while also being present throughout most months in both years (Figure 2). May of Year 1 was dominated by *C. harengus* and *P. virens* reads, whereas May in Year 2 was dominated by *S. sprattus* reads. *Ctenolabrus rupestris* reads dominated July of Year 1, whereas in July of Year 2 dominance was shared by *Ctenolabrus rupestris*, *Platichthys flesus* and *G. flavescens*. September of Year 1 was dominated by *S. scombrus* and *G. flavescens* reads, while September of Year 2 was dominated by *G. flavescens* reads. November of both years was dominated by *G. flavescens* reads. January of Year 1 shows similar read abundance for several species, without any clear dominant species, while January of Year 2 was dominated by *S. scombrus*.

For the MiFish primer set, the Friedman test showed a significant difference in the total read abundance between months within Year 1 (χ2 = 13.26, df = 5, p = 0.021), and between months within Year 2 (χ2 = 37.91605, df = 5, p = 3.923e-07). The Friedman test also indicated a significant difference between months within Year 1 and within Year 2 of the Elas02 primer set, (χ2 = 34.649, df = 5, p= 1.77e-06) and (χ2 = 73.3005, df = 5, 2.10e-14) respectively.

We found seasonal read count variations for teleost species detected by the MiFish primer set across both sampled years (Year 1: reanalyzed data from Carvalho et al. (2024); Year 2: newly collected data) (Figure 2). Most species, including *Clupea harengus* (Atlantic herring) and *Sprattus sprattus* (European sprat), were consistently detected in both years. *Clupea harengus* exhibited peak read counts in March, while *Sprattus sprattus* peaked in May. Read counts for the majority of species remained stable across months and years. All species detected in Year 1 were also identified in Year 2, though 19 additional taxa—including *Abramis brama* (common bream), *Anguilla anguilla* (European eel), *Melanogrammus aeglefinus* (haddock), and *Trachurus trachurus* (Atlantic horse mackerel)—were uniquely detected in Year 2.

Using the Elas02 primer set, *Clupea harengus* and *Sprattus sprattus* showed consistent peaks in March and May, respectively (Figure 3). Most species were detected in both years, though a subset differed between sampling periods: *Amblyraja radiata* (thorny skate), *Centrolabrus exoletus* (rock cook), *Chimaera monstrosa* (rabbitfish), and *Zoarces viviparus* (viviparous blenny) were unique to Year 1, while *Abramis brama*, *Alburnus alburnus* (common bleak), *Trisopterus esmarkii* (Norway pout), and *Zeugopterus norvegicus* (Norwegian topknot) were exclusive to Year 2. Read counts for dominant taxa remained stable across months, mirroring the results observed with MiFish.

**Figure 3.**
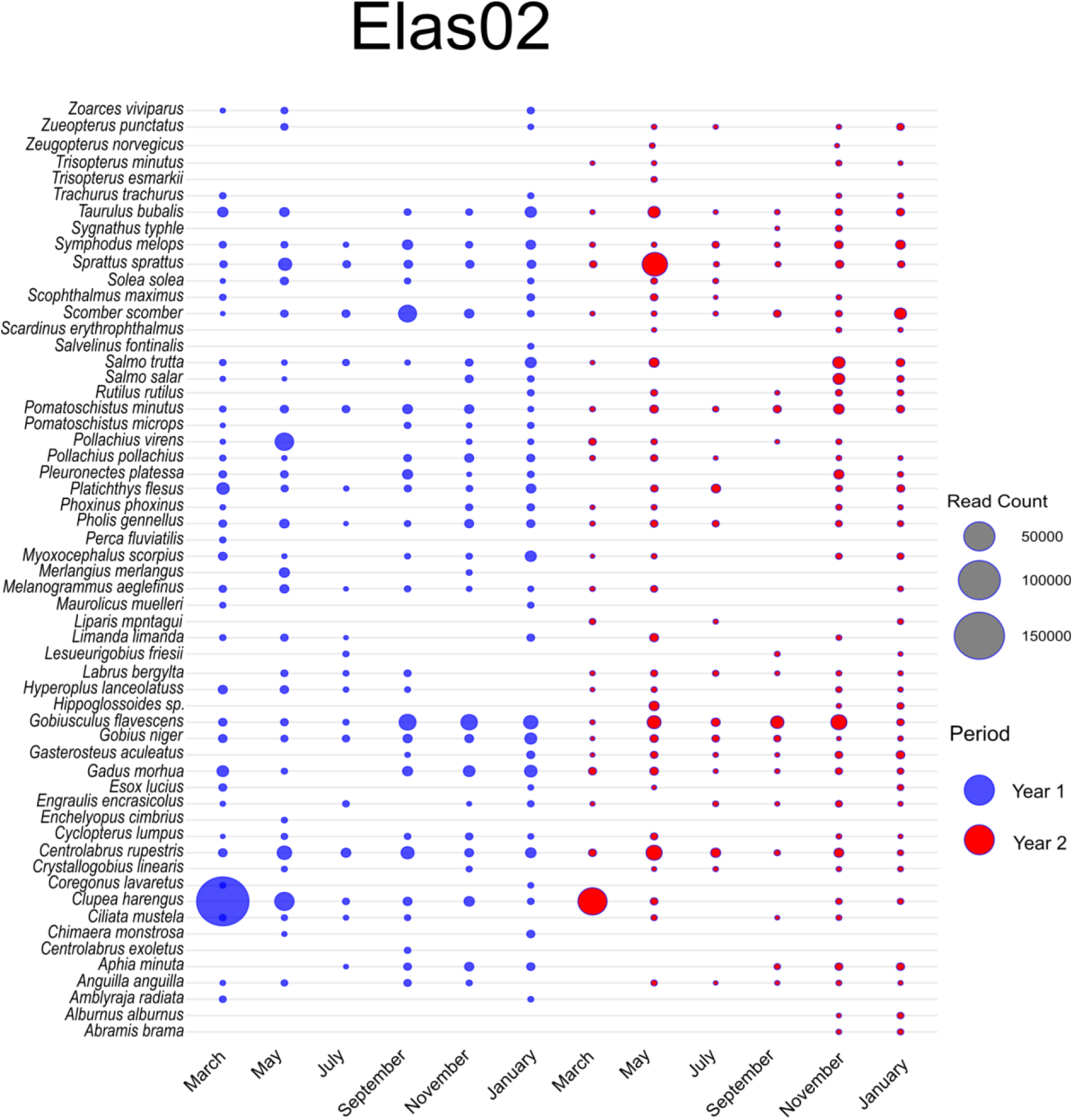
Bubble plot showing the relative read abundance of each detected fish species using the Elas02 primer set across all sampling events

The recovered species richness differed between primers and years (Figure 4). The MiFish primer detected significantly higher richness in Year 2 compared to Year 1 (Wilcoxon rank sum test, *p* = 0.000492), with January exhibiting the highest richness in both years (Figures 4A, C). In contrast, the Elas02 primer showed no significant interannual difference in richness (Figures 4B, D), though January similarly hosted the highest species counts for both years. Notably, species detected in Year 2 but undetected in Year 1 using MiFish (e.g., *Melanogrammus aeglefinus*, *Trachurus trachurus*) were detected in both years using Elas02, highlighting that the lower fish read count for MiFish in Year 1 is associated with a fish diversity loss. While *Anguilla anguilla* was detected by both primers in Year 2, Elas02’s richness remained stable across years despite turnover in specific taxa (e.g., loss of *Amblyraja radiata* in Year 2) (Figure 2 and 3).

**Figure 4.**
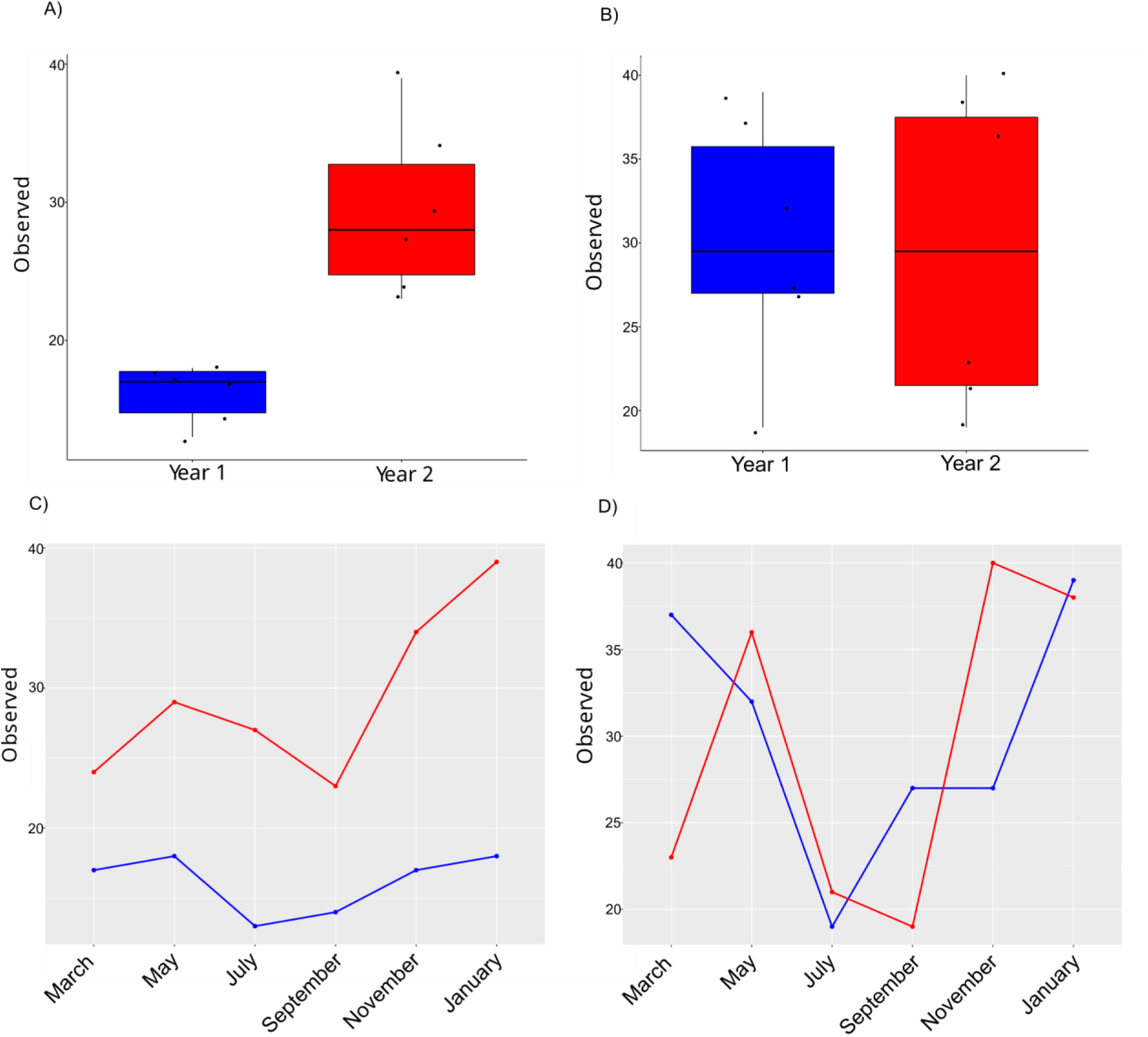
Variation of the species richness observed with both MiFish and Elas02 primer sets across the two sampled years

We found that the number of fish species retrieved using the MiFish primer set was highly variable across both years, with 19 fish species detected in Year 1 and 44 detected in Year 2. However, using the Elas02 primer set, the number of species was similar in both years, with 48 and 46 fish species detected during Year 1 and Year 2, respectively (Figures 1 and 4).

NMDS ordination based on Bray-Curtis dissimilarity revealed partial separation of Year 1 and Year 2 samples across both primers (MiFish and Elas02), with samples from the same year generally aligning on the same side of the ordination space for rarefied and presence/absence datasets (See Zenodo repository at https://doi.org/10.5281/zenodo.14809757). PERMANOVA results indicated contrasting patterns between data types, and for rarefied datasets, no significant compositional differences were detected between years (MiFish: *p* > 0.05; Elas02: *p* > 0.05). In contrast, presence/absence transformations showed statistically distinct communities between years for both primers (MiFish: *p* = 0.002; Elas02: *p* = 0.021).

Some of the fish species initially identified through the taxonomic assignment using the GBIF-mediated MitoFish database led to false positive detections of 13 species, which are unknown to occur in the Oslofjord area or all of Norway and have not been detected in similar studies of the Oslofjord area (Carvalho et al., 2024, Kvalheim et al., 2024). By comparing the DNA sequences of these 13 unexpected fish species to the GenBank database using BLAST, 12 of them could be taxonomically re-assigned to fish species known to occur in the area with records of observation and/or detection in Artsdatabanken and the GBIF database and with high identity percentage values to the reference sequences in GenBank. More specifically, *Chelidonychthys kumu* was re-assigned to *Chelidonichthys lucerna*, *Clupea pallasii* to *Sprattus sprattus*, *Hippoglossoides elassodon* to *Hippoglossoides sp.*, *Lepidorhombus whiffiagonis* to *Zeugopterus norvegicus*, *Limanda aspera* to *Limanda limanda*, *Liopsetta glacialis* to *Platichthys flesus*, *Liparis agassizii* to *Liparis montagui*, *Myoxocephalus thompsonii* to *Myoxocephalus scorpius*, *Pleuronectes quadrituberculatus* to *Pleuronectes platessa*, *Salmo labrax* to *Salmo trutta*, *Symphodus ocellatus* to *Symphodus melops*, and *Zoarces americanus* to *Zoarces viviparus*. All of these 12 re-assigned fish species, except for *Zeugopterus norvegicus*, have also been detected using eDNA by Carvalho et al. (2024) or have been previously observed in Norway (Kvalheim et al., 2024). The DNA sequence assigned to *Collichthys lucidus*, which was detected once during year 2 with the MiFish primer set, could not be re-assigned to any fish species with records of occurrence or observation in the Oslofjord area or in all of Norway, and could be resulting from human consumption through imported food items.

Like *Zeugopterus norvegicus*, there are four other fish species that have been identified in this study, which were not detected or observed by Carvalho et al. (2024) and Kvalheim et al. (2024) but have occurrence or observation records in Artsdatabanken and the GBIF databases: *Hyperoplus lanceolatus*, *Merluccius merluccius*, *Alburnus alburnus*, and *Abramis brama*. *H. lanceolatus* was found during multiple sampling events in both years using both primer sets, while *M. merluccius* was detected at two events in Year 2 by just the MiFish primer set. *A. alburnus* and *A. brama* were detected by both primer sets, but only at two and three sampling events during Year 2, potentially due to particular winter conditions leading these to find refuge in the fjord or to DNA transport from rivers.

## Discussion

Here, we used eDNA metabarcoding to analyse fish communities over two years using the MiFish and Elas02 primer sets, both regularly used in fish metabarcoding studies (Burian et al., 2023; Maiello et al., 2024; Miya et al., 2015; Ward et al., 2022), and found significant differences in species composition and relative abundance between sampled years and sampling events. We built on the foundational work from Carvalho et al. (2024) initially assessing the use of eDNA detection to explore fish assemblage in the Oslofjord. In addition to the mobilization and reanalysis of their raw sequencing data corresponding to the first year of the survey, we analysed eDNA samples for an additional year using the same primer sets. Using a different bioinformatic pipeline and reference database linked to the taxonomic assignment, we clarified key patterns and uncovered new insights into how fish communities change across seasons and years. For example, both studies agreed on one clear trend: *Clupea harengus* dominated March populations in both years, regardless of the primers used. This consistency suggests *C. harengus* is a reliable indicator of early spring conditions in the ecosystem (Carvalho et al., 2024; Dzadey, 2014). Autumn months (September–November) showed strong similarities, with *S. scombrus* (Scombridae) and *G. flavescens* (Gobidae) displaying high read abundance as in (Carvalho et al., 2024), suggesting that these species define late-season assemblages. Following the reanalysis of the first year sampling data, we found no clear dominant species in January 2023, but identified *S. scombrus* as a dominant species in January 2024 using the MiFish primer set, highlighting how extended sampling can reveal dynamic changes single-year studies might miss (Courtaillac et al., 2024; Zinger et al., 2019).

The repeated detection of *C. harengus* and *S. sprattus* peaks in March and May (Figure 2 and 3) across both primers and years highlights their consistent role as seasonal markers, likely tied to predictable behaviors like spawning or migration (Bekkevold et al., 2005; Cushing, 1969). The new species detected in Year 2 using MiFish primer set—such as *Anguilla anguilla* (European eel) and *Melanogrammus aeglefinus* (haddock)—could reflect actual changes in the ecosystem between years (e.g., shifts in habitat use or population size) or simply the expanded opportunity to detect rare species with an additional year of sampling (Stokoele et al., 2020). Indeed, migration phenology can change, and the peak of migration could have been missed in Year 1 but observed in Year 2. However, it should be noted that the lowest fish species richness retrieved using the MiFish primer set during Year 1 could be an effect of the lower read count obtained compared to Year 2. Such lower output compared to the following year or Elas02 primer set could be due to library preparation or sequencing issues. As a result, it is unsure if the newly detected species in Year 2 using the MiFish primer set reflect biological or seasonal events or are linked to analytical challenges in Year 1. Similarly, the absence of species like *Amblyraja radiata* (thorny skate) in Year 2, despite Elas02 primer set detecting them in Year 1, reminds us that low-abundance marine life can be inconsistently captured by eDNA, even with targeted primers (Ficetola et al., 2015). While our methodological variation in data analysis (e.g., size filtering to reduce non-target DNA) clarified some patterns—such as resolving *S. scomber* as stable winter residents—the core strength of this work lies in its two-year perspective. By pairing Carvalho et al. (2024) foundational data with our follow-up sampling, we demonstrate that multi-year eDNA research may separate transitory findings from long-term ecological trends without overstating technological gains.

Differences between Year 1 and Year 2 were potentially caused by species turnover (presence/absence changes) rather than abundance changes. Rarefied data, which retained abundance information but standardised sequencing depth, showed no significant differences, indicating dominant species’ abundances remained stable. Presence/absence transformations, however, highlighted turnover—likely from rare or transient species— emphasise their sensitivity to low-frequency taxa. This divergence likely emphasises that data choice (abundance vs. incidence) influences ecological interpretation: turnover may reflect environmental changes or dispersal dynamics overlooked by abundance-focused approaches (Fonseca 2018). While results were consistent across primers, longer-term data is needed to understand whether these changes represent stochastic variation or consistent trends (Mathieu et al., 2020; Zinger et al., 2019). Together, these findings show how combining approaches—learning from previous studies while refining methods—helps separate true ecological trends from methodological noise. Even while no single study can fully represent the complexity of nature, combining several approaches and years of research helps us better understand how fish groups adapt to a changing environment.

Overall, our findings also concur with the results highlighted by Kvalheim et al. (2024) and regarding the use of eDNA metabarcoding as a tool for fish monitoring in the Oslofjord, but further highlights its benefits using time series analysis. Kvalheim et al. (2024) combined citizen science sampling with eDNA metabarcoding analysis of 96 sites using the MiFish primer set (Miya et al. 2015) to characterise the Oslofjord’s fish biodiversity over the summer of 2022. In addition, the study combined DADA2 as a bioinformatic pipeline, and taxonomic assignment using the Basic Local Alignment Search Tool (BLAST) in conjunction with an inhouse fish database, ScandiFish v. 1.4 (Stensrud, 2022). We detected similar fish species (61 in this study and 57 in Kvalheim et al 2024). Freshwater species known to be present in Norway but undetected using eDNA in Kvalheim et al. (2024) were detected in our study, such as *Esox lucius*, *Rutilus rutilus* and *Scardinius erythrophthalmus*. Furthermore, *E. lucius,* together with other species such as *Trachurus trachurus* and *Trisopterus esmarkii* were detected by the Elas02 primer sets in our study, but not in (Kvalheim et al., 2024), highlighting the benefits of using multiple primer sets to optimise species detection (Burian et al., 2022).

Finally, it should be noted that methodological decisions have strong impacts on the final results. Here, despite using different sampling protocols, bioinformatic tools and reference databases, we detected similar species in (Kvalheim et al., 2024) and (Carvalho et al., 2024). However, the initial species list obtained following taxonomic assignment using the pipeline and tools provided by GBIF relied on the MitoFish v3.97 database, resulting in the false positive detection of 13 different species (Iwasaki et al 2013; Sato et al 2018). This highlights the importance of using local reference databases, and the necessity to validate species detection using previous species records, in this case through GBIF and Artsdatabanken databases. Indeed, despite using reliable and curated reference databases, species identification of metabarcoding data relies on the taxonomic assignment of short amplicons, potentially leading to issues regarding the taxonomic resolution when identifying closely related species (Burian et al., 2023; Polanco et al., 2021; Taberlet et al., 2012).

## Conclusion

Our two-year eDNA metabarcoding study of Oslofjord revealed dynamic shifts in fish community composition, with seasonal dominance of key species and interannual turnover of rare taxa. We observed variation in communities between years, underlining the importance of species presence/absence shifts in explaining abundance fluctuations. The detection of additional species in the second sampled year revealed the sensitivity of eDNA in capturing ecological variability together with time series analysis. These results support the use of long-term monitoring to differentiate transient signals from permanent trends, particularly in ecosystems subject to anthropogenic stress. This study suggests that eDNA based monitoring is a reliable method for assessing marine biodiversity and informs adaptive management techniques in rapidly changing marine habitats by combining temporal resolution with primer-specific insights.

## Supporting information

All supplementary figures, tables and files

## Acknowledgments

We would like to thank the Biological Marine station in Drøbak for the help and support regarding sample collection over the last years. We thank the DNA lab at the Natural History Museum, UiO, and especially Jarl Andreas Anmarkrud and Birgitte Lisbeth Graae Thorbek for their support and assistance.

## Authors Contribution

Conceptualization: QM, HdB; Sampling: SM, IE, COC, QM; Laboratory analysis: SM, COC, ASN; Bioinformatics: QM, SM, COC; Data curation: QM, IE, SM; Statistical analysis: SM; Original draft: SM, IE, COC, QM; Review and editing: all authors; Funding acquisition: QM, MD, HEK, HdB.

## Data availability statement

The raw sequencing data that support the findings of this study, bioinformatic scripts, mapping files and metadata are openly available in Zenodo at https://doi.org/10.5281/zenodo.14809757.

## Supplementary information

Table S1. Information about the eDNA samples collected in the second year Table S2. Species presence across primers and years

Figure S1. NMDS plots using rarefied and presence/absence data

