## Supplementary figures and images for "Investigating the dynamics of the aquatic community in Oslofjord through time series analysis of eDNA"

### Figure_S1.pdf

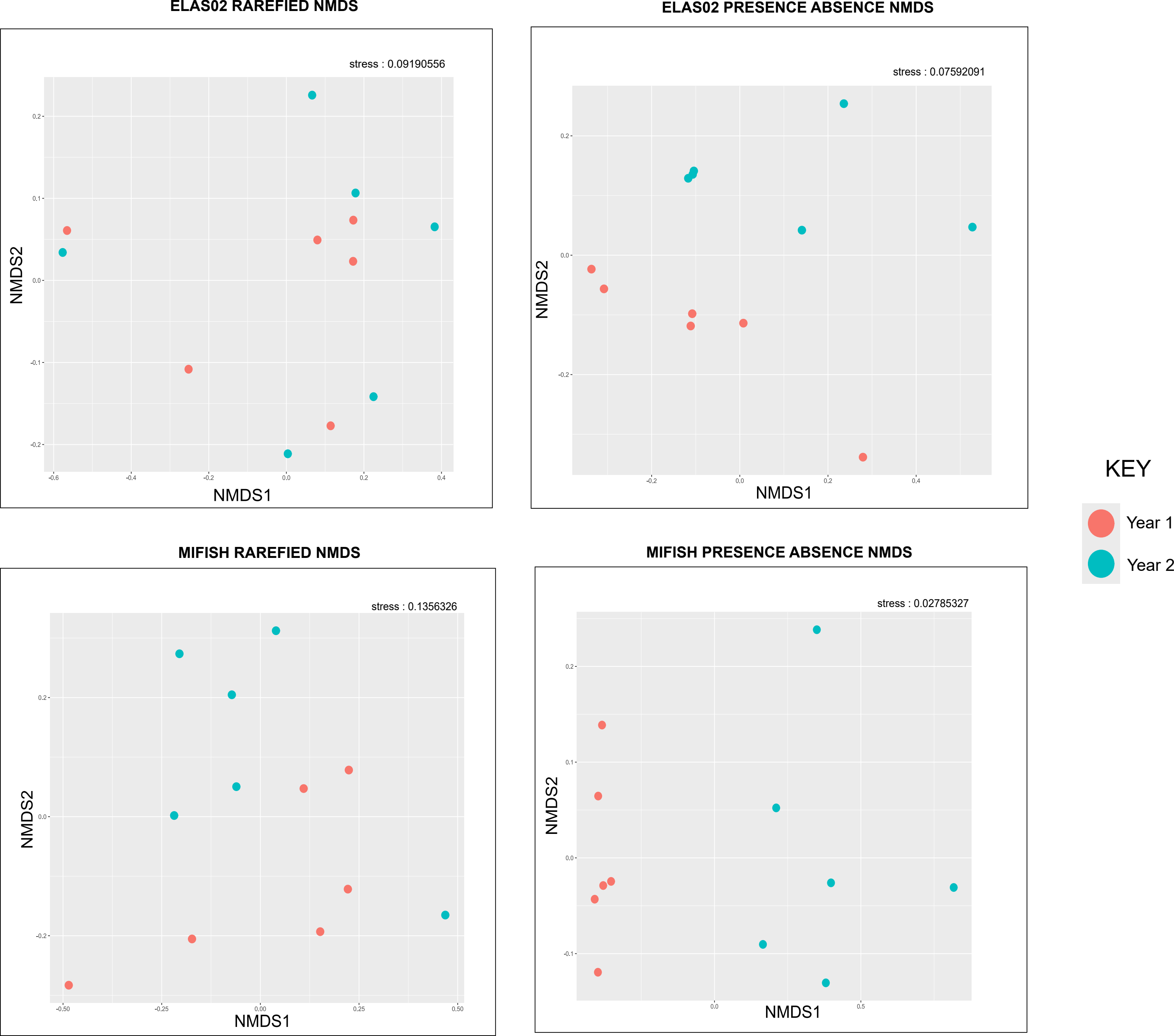
